# Metabolic effects of risk alleles in *PNPLA3, TM6SF2, GCKR* and *LYPLAL1* inform about heterogeneity of non-alcoholic fatty liver disease

**DOI:** 10.1101/143511

**Authors:** Eeva Sliz, Sylvain Sebert, Peter Würtz, Antti J Kangas, Pasi Soininen, Terho Lehtimäki, Mika Kähönen, Jorma Viikari, Minna Männikkö, Mika Ala-Korpela, Olli T Raitakari, Johannes Kettunen

## Abstract

Fatty liver has been associated with unfavourable metabolic changes in circulation and is considered as a risk factor for cardiometabolic complications such as type 2 diabetes and cardiovascular disease. We aimed to provide insights in fatty liver related metabolic deviations by studying the resemblance between the metabolic profile associated with fatty liver observationally and metabolic profiles of non-alcoholic fatty liver disease (NAFLD) risk increasing genotypes. We determined cross-sectional associations of ultrasound-ascertained fatty liver status with 123 metabolic traits in 1,810 individuals aged 34-49 years from The Cardiovascular Risk in Young Finns Study. The cross-sectional associations were compared with the association profiles of NAFLD risk alleles in *PNPLA3*, *TM6SF2*, *GCKR*, and *LYPLAL1* with the corresponding metabolic traits obtained from a publicly available genome-wide association study including up to 24,925 European individuals. The analysis revealed substantially different metabolic effects of the risk alleles. *PNPLA3* rs738409-G, the strongest genetic risk factor to NAFLD, did not associate with metabolic changes. *GCKR* rs1260326-T resulted in an association profile similar to the observational fatty liver associations. Metabolic effects of *LYPLAL1* rs12137855-C were similar, but statistically less robust, to the effects of *GCKR* rs1260326-T. In contrast, NAFLD risk allele *TM6SF2* rs58542926-T displayed opposite metabolic associations when compared with the observational association pattern.

**Conclusion:** The divergent effects of the risk alleles on circulating lipids and metabolites underline involvement of several metabolic pathways in NAFLD and suggest that there are pathogenically different subtypes of NAFLD with alternate metabolic consequences. NAFLD risk alleles may have neutral or even cardioprotective effect on circulating lipids and metabolites providing evidence that hepatic lipid accumulation by itself would not necessarily cause the metabolic deviations associated observationally with fatty liver.

---

Non-alcoholic fatty liver disease (NAFLD) covers a range of liver disorders that originate from excessive lipid, mainly triglyceride, accumulation in the liver (1). Fatty liver has been proposed to result in dyslipidemia due to increased secretion of very low-density lipoproteins (VLDL) and impaired clearance of intermediate and low-density lipoproteins (IDL, LDL) from circulation (2, 3). Studies using detailed metabolic profiling have shown that fatty liver associates with a wide range of metabolic aberrations in circulation (4–6). Kaikkonen *et al.* assessed cross-sectional and prospective associations of fatty liver with 68 circulating metabolic measures in the population-based Young Finns Study (YFS): the most prominent associations were seen with triglycerides in the largest VLDL particles as well as with VLDL particle concentrations, but fatty liver was robustly associated also with many non-lipid traits, such as branched-chain amino acids leucine and isoleucine (4). Observations from animal models and human studies have shown that liver fat content correlates with levels of blood lipids and glucose (7–9). However, fatty liver does not always seem to relate to dyslipidemia or changes in glycemic traits and thus its effects on metabolic changes in circulation has remained unclear (10, 11).

Genetic factors contribute to the pathogenesis of fatty liver. Four DNA sequence variants, Patatin-like phospholipase domain containing 3 (*PNPLA3*) rs738409-G, Glucokinase regulator (*GCKR*) rs780094-T, Neurocan (*NCAN*) rs2228603-T, and Lysophospholipase-like 1 (*LYPLAL1*) rs12137855-C, were associated with computed tomography defined steatosis and biopsy-proven NAFLD involving lobular inflammation and fibrosis in a large-scale genome-wide association study (GWAS) (12). Variants in these four loci have also been used to determine a genetic risk score for NAFLD (13). Other studies have shown that *GCKR* rs1260326-T and Transmembrane 6 superfamily member 2 (*TM6SF2*) rs58542926-T are the functional variants at the *GCKR* and *NCAN* loci, respectively (14–16). *PNPLA3* rs738409-G is the strongest genetic risk factor for NAFLD (17) having an odds ratio of 3.24 for histologic NAFLD (12). A meta-analysis on 2,937 individuals with biopsy-diagnosed NAFLD showed that the GG genotype increases hepatic triglyceride content by 73% in comparison to reference CC genotype (18). *GCKR* rs1260326-T and *TM6SF2* rs58542926-T are also recognized as important determinants of inter-individual variation in liver fat (14, 19, 20). Among the aforementioned variants, the function of *LYPLAL1* rs12137855-C is the least known.

In the present study we perform detailed metabolic profiling of fatty liver in young and middle-aged adults with ultrasound ascertained fatty liver. To add insights into molecular mechanisms of fatty liver related metabolic aberrations, we compare how the observational fatty liver associations match with metabolic association profiles of the aforementioned NAFLD risk alleles. We utilize publicly available summary statistics from a metabolomics GWAS to assess the detailed metabolic effects of the risk variants (21). Understanding the relation between fatty liver and changes in circulating lipids and metabolites is helpful in acquiring more opportunities for treatment and prevention of this complex condition and related cardiometabolic complications.

## Methods

### Cross-sectional fatty liver associations with circulating metabolites

The Cardiovascular risk in Young Finns Study (YFS) is a population based follow-up study started in 1980. In 2011, 2,046 individuals aged 34 to 49 years participated to ultrasound imaging (Acuson Sequoia 512, Acuson, Mountain View, CA, USA) of the liver. A trained sonographer assessed the presence of fatty liver using 4.0 MHz adult abdominal transducers, and the participants were categorised into two groups: fatty liver and no fatty liver. In total, fatty liver was diagnosed in 18.6% (N=372) of the participants. The population and ultrasound imaging of the liver is described in more detail in (4). After exclusion of pregnant women and individuals using lipid lowering medication or oral contraceptives, the total number of individuals included to determine the associations between fatty liver and metabolic phenotypes was 1,810 (N_fatty liver_=338). The study was approved by the local Ethics Committee and written informed consent was obtained from all the participants.

### NAFLD risk allele associations with circulating metabolites

Effect estimates of *PNPLA3* rs738409-G, *GCKR* rs1260326-T, *GCKR* rs780094-T, *TM6SF2* rs58542926-T, *NCAN* rs2228603-T, and *LYPLAL1* rs12137855-C on the metabolic traits were acquired from a published GWAS performed using 14 European cohorts from Finland, Germany, the Netherlands and Estonia adding up to 24,925 individuals (21). The mean age and BMI of the participants per cohort ranged from 23.9 to 61.3 years and from 23.1 to 28.2 kg/m^2^ with the whole sample means being 46.3 years and 26.0 kg/m^2^.

Risk allele frequencies and odds ratios to NAFLD for the studied loci are described in Table 1. To facilitate comparison of the risk allele associated metabolic changes relative to NAFLD risk increase, the risk allele effects and corresponding standard errors were scaled with respect to the log(odds ratio) on histologic NAFLD associated with the corresponding locus in a large-scale GWAS (12).

**Table 1.**
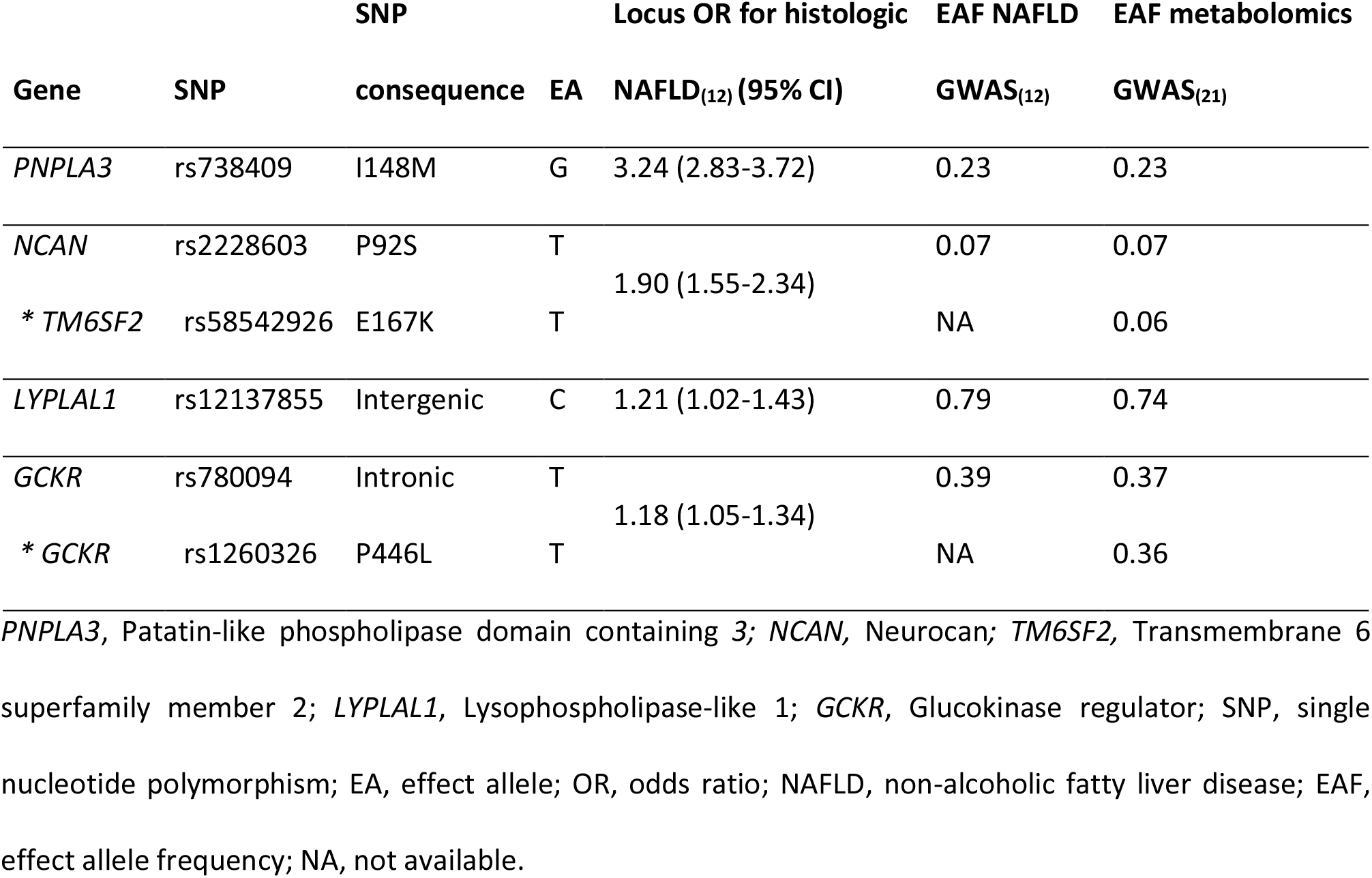
Description of NAFLD risk increasing genotypes extracted from the open access data. DNA sequence variants studied in the present study were associated with computed tomography characterized steatosis and biopsy-proven NAFLD involving liver inflammation and fibrosis by Speliotes *et al*. (12). The functional variants explaining the NAFLD associations of *NCAN* rs2228603 and *GCKR* rs780094 are denoted separately *. To achieve coherency, the *NCAN* locus is referred as *TM6SF2* throughout the paper.

### Metabolic profiling

The metabolic profiles of all samples included were assessed using a nuclear magnetic resonance (NMR) metabolomics platform (Nightingale Health, Helsinki, Finland) described in (21, 22) and reviewed in more detail in (23, 24). The platform covers multiple distinct metabolic traits including lipoprotein subclasses and their lipids, fatty acids, amino acids and glycolysis precursors. To determine the cross-sectional metabolic aberrations associated with fatty liver, we utilized 123 metabolic measures representing a broad molecular signature of systemic metabolism (Supplementary Table 1).

### Statistical analyses

All analyses were done using R version 3.2.2. Statistical significance was considered at P<0.002 (0.05/22), where 22 is the number of principal components explaining 95% of the variation in the NMR metabolomics data (21).

#### Cross-sectional associations

Linear regression models were fitted to determine the cross-sectional associations of fatty liver with each of the metabolic measures in the YFS population. To facilitate the comparison of observed and genetic effect estimates, the data processing and analysis model were done correspondingly to Kettunen *et al.* (21): the metabolic phenotypes were adjusted for age, sex, and ten first genetic principal components preceding the analysis, and the resulting residuals were transformed to normal distribution by inverse rank-based normal transformation. The adjusted and transformed metabolic phenotypes were used as outcomes in the equations and fatty liver served as a categorical variable (fatty liver vs. no fatty liver).

#### Comparisons of the metabolic association profiles

The overall resemblance between the risk alleles and observational fatty liver effects on metabolites was determined by a linear fit of each of the risk allele association profile versus cross-sectional fatty liver association profile (25, 26). In addition, the linear fit of all the pairs of the risk allele association profiles were determined in order to identify alleles with similar metabolic effects informing about gene products functioning in related biological pathways. This method was also used to ensure that the metabolic profiles of the NAFLD risk alleles *GCKR* rs780094-T and *NCAN* rs2228603-T correspond to ones of *GCKR* rs1260326-T and *TM6SF2* rs58542926-T that have been identified as the functional variants in *GCKR* and *NCAN/TM6SF2* loci, respectively (14–16).

## Results

### Fatty liver and circulating metabolites

Characteristics of the YFS study population are shown in Table 2. Fatty liver was associated with 88 metabolic phenotypes (P<0.002) in the YFS population when adjusted with age, sex, and ten genetic principal components (Panel 1 in Figure 1; Supplementary Figure 1; Supplementary Table 2). Lipoprotein subclass concentrations showed a trend where the largest VLDL particles displayed the most pronounced associations that got weaker while the particle size decreased, and again stronger for medium and small LDL particles (Panel 1 in Figure 1). Concentration of very large and large HDL particles showed negative association with fatty liver, while small HDL particles displayed a strong positive association. Fatty liver was also associated with increased VLDL diameter while the associations with LDL and HDL particle diameters were negative. Fatty liver associated negatively with apolipoprotein A-I and positively with apolipoprotein B. Concentrations of serum triglycerides and triglycerides in all lipoprotein subclasses were increased. Fatty liver was also associated with increased level of saturation of circulating fatty acids along with increased concentrations of circulating fatty acids, while it showed a negative association with fatty acid length. In addition, fatty liver displayed positive association with glucose, lactate, pyruvate, and glycerol, as well as with amino acids alanine, isoleucine, leucine, valine, phenylalanine, and tyrosine. Association between fatty liver and amino acid glutamine was negative.

**Figure 1.**
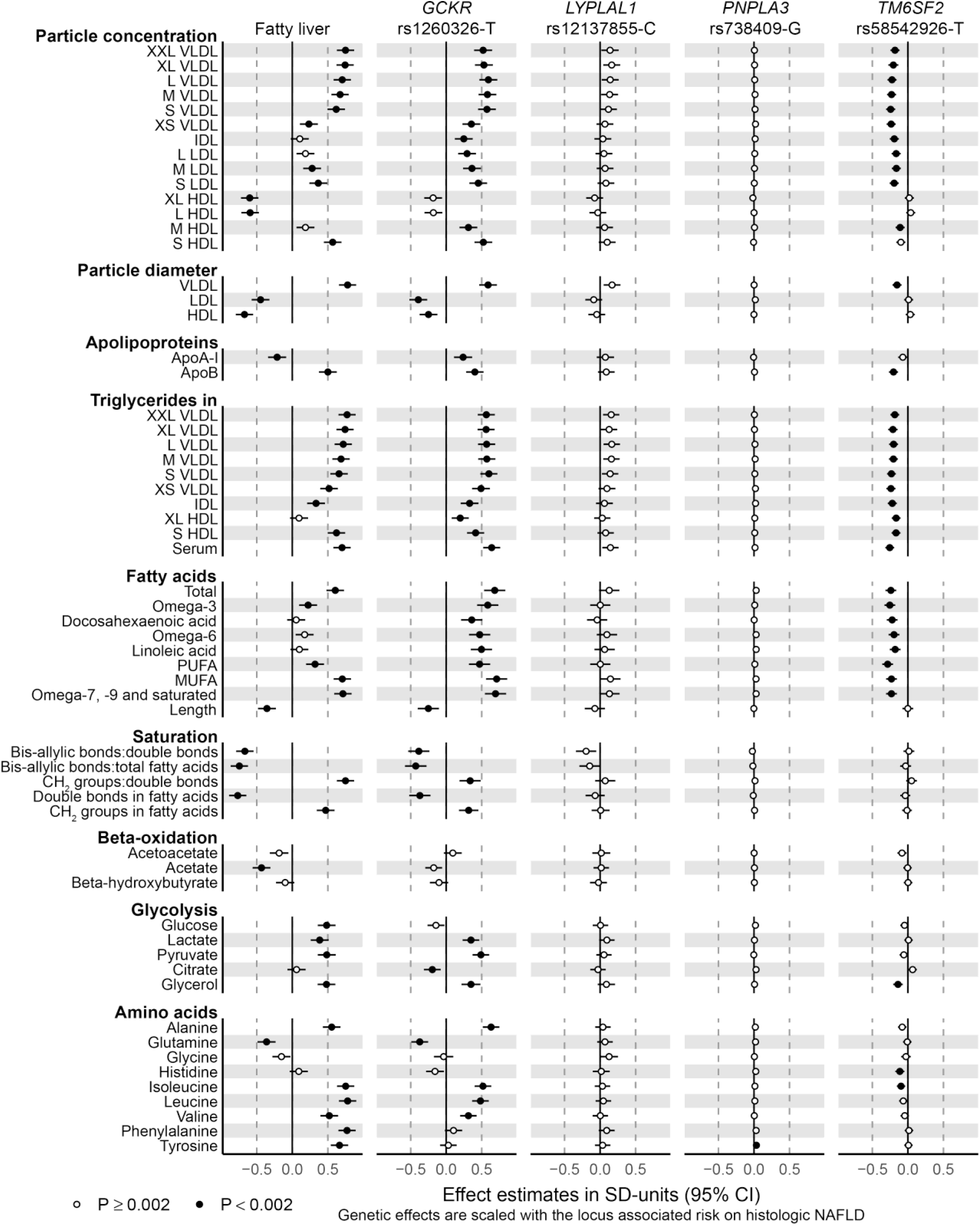
Cross-sectional associations of fatty liver with lipoprotein particle subfraction concentrations, lipoprotein particle diameter, apolipoproteins, triglycerides, fatty acids, fatty acid saturation, beta-oxidation, glycolysis and amino acid related metabolites, and the corresponding associations with four NAFLD risk alleles. Cross-sectional associations were determined in 1,810 adults aged 34-49 years of whom 338 were diagnosed with ultrasound-based fatty liver. The metabolic phenotypes were adjusted for age, sex, and ten first genetic principal components prior to analysis. Genetic effects of the NAFLD risk alleles *GCKR* rs1260326-T, *LYPLAL1* rs12137855-C, *PNPLA3* rs738409-G and *TM6SF2* rs58542926-T were acquired from a metabolomics GWAS including up to 24,925 Europeans (21). Genetic effect estimates were scaled with respect to the NAFLD risk associated with the corresponding locus (12).

**Table 2.** Study population: YFS.

|  | **Fatty liver** | **No fatty liver** | **Total population** |
| --- | --- | --- | --- |
| **N** | 338 | 1,472 | 1,810 |
| **Male (%)** | 237 (70.1) | 630 (42.8) | 867 (47.9) |
| **BMI, kg/m<sup>2</sup>** | 30.5 ± 5.3 | 25.5 ± 4.3 | 26.4 ± 4.9 |
| **Age in years** | 43.0 ± 4.7 | 41.6 ± 5.0 | 41.9 ± 5.0 |
| **Smokers (%)** | 55 (16.3) | 198 (13.5) | 253 (14.0) |
| **Alcohol consumption per day\*** | 2.56 ± 1.7 | 2.09 ± 1.5 | 2.18 ± 1.5 |

\* One unit equals to 12 g of alcohol.
Values are mean ± SD, or absolute N count and corresponding percentage.
YFS, Young Finns Study.

### NAFLD risk alleles in *GCKR* and *LYPLAL1* tend to increase concentrations of circulating lipids

The studied NAFLD risk increasing alleles showed different association profiles on metabolic measures. *GCKR* rs1260326-T associated with increased particle concentrations of all VLDL, IDL, and LDL subclasses as well as concentrations of medium and small HDL particles (Panel 2 in Figure 1; Supplementary Figure 2; Supplementary Table 3). The amount of triglycerides was increased in all the lipoprotein subclasses. *GCKR* rs1260326-T was associated with increased diameter of VLDL particles, while the association with LDL and HDL particle diameters were negative. The variant showed positive association with concentrations of apolipoproteins A-I and B, as well as with all the studied fatty acids, and displayed negative association with fatty acid length. *GCKR* rs1260326-T associated also with increased fatty acid saturation. *GCKR* rs1260326-T was associated positively with glycolysis related metabolites lactate, pyruvate, and glycerol, while association with citrate was negative. This risk allele associated positively with amino acids alanine, isoleucine, leucine, and valine, and negatively with glutamine. The results of *GCKR* rs780094-T, a variant originally identified as a NAFLD risk allele in the GCKR locus (12) being in linkage disequilibrium with the functional *GCKR* rs1260326-T (14), were highly similar to the results of *GCKR* rs1260326-T and can be found in the supplementary material (Supplementary Figure 1; Supplementary Figure 2; Supplementary Table 3).

The metabolic association profile of *LYPLAL* rs12137855-C was highly similar to the association profile of *GCKR* rs1260326-T in terms of highly correlated point estimates of the two risk alleles (Supplementary Figure 3A), but the effects of *LYPLAL* rs12137855-C were statistically less robust and the effect magnitudes were weaker than for the *GCKR* variant (Panel 3 in Figure 1; Supplementary Figure 2; Supplementary Table 3).

### *PNPLA3* rs738409-G does not show association with circulating metabolic traits

*PNPLA3* rs738409-G, the strongest genetic contributor to the hepatic fat content (17), displayed metabolic association profile close to null (Panel 4 in Figure 1; Supplementary Figure 2; Supplementary Table 3). When compared with the cross-sectional fatty liver effects, the NAFLD-scaled metabolic effects of *PNPLA3* rs738409-G were much closer to zero, and the confidence intervals for effect estimates of cross-sectional fatty liver and *PNPLA3* rs738409-G were clearly separated.

### *TM6SF2* rs58542926-T associates with lower-risk metabolic profile

*TM6SF2* rs58542926-T displayed strong association profile throughout the studied metabolites (Panel 5 in Figure 1; Supplementary Figure 2; Supplementary Table 3). However, the associations were negative indicating reduced concentrations of the lipids and metabolites in relation to higher NAFLD risk. The *TM6SF2* variant was observed to decrease concentrations of all the VLDL, IDL and LDL particle subclasses and all the lipid species in these subclasses. In addition, the variant associated inversely with serum total triglycerides and triglycerides in all the lipoprotein subclasses including the HDL subclasses. The diameter of the VLDL particles was reduced, while the variant did not contribute to the LDL nor HDL particle diameters. The *TM6SF2* rs58542926-T associated also with decreased concentration of the apolipoprotein B but did not influence concentration of apolipoprotein A-I. It associated inversely also with all the fatty acid concentrations, while the qualitative measures such as fatty acid length or saturation measures were not influenced by this variant. In addition, concentrations of histidine, isoleucine and glycerol were reduced. The results for *NCAN* rs2228603-T in the same NAFLD risk locus can be found in the supplement (Supplementary Figure 1; Supplementary Figure 2; Supplementary Table 3).

### Resemblance of the metabolic effects

The correspondence between the metabolic effects of the risk alleles and observational fatty liver was the highest between the *GCKR* rs1260326-T and fatty liver (R^2^ = 0.77; Figure 2C). The remaining coefficients of determination were R^2^ = 0.67 for *LYPLAL1* rs12137855-C versus fatty liver (Figure 2D), R^2^ = 0.45 for *TM6SF2* rs58542926-T versus fatty liver (Figure 2B), and R^2^ = 0.30 for *PNPLA3* rs738409 G versus fatty liver (Figure 2A).

**Figure 2.**
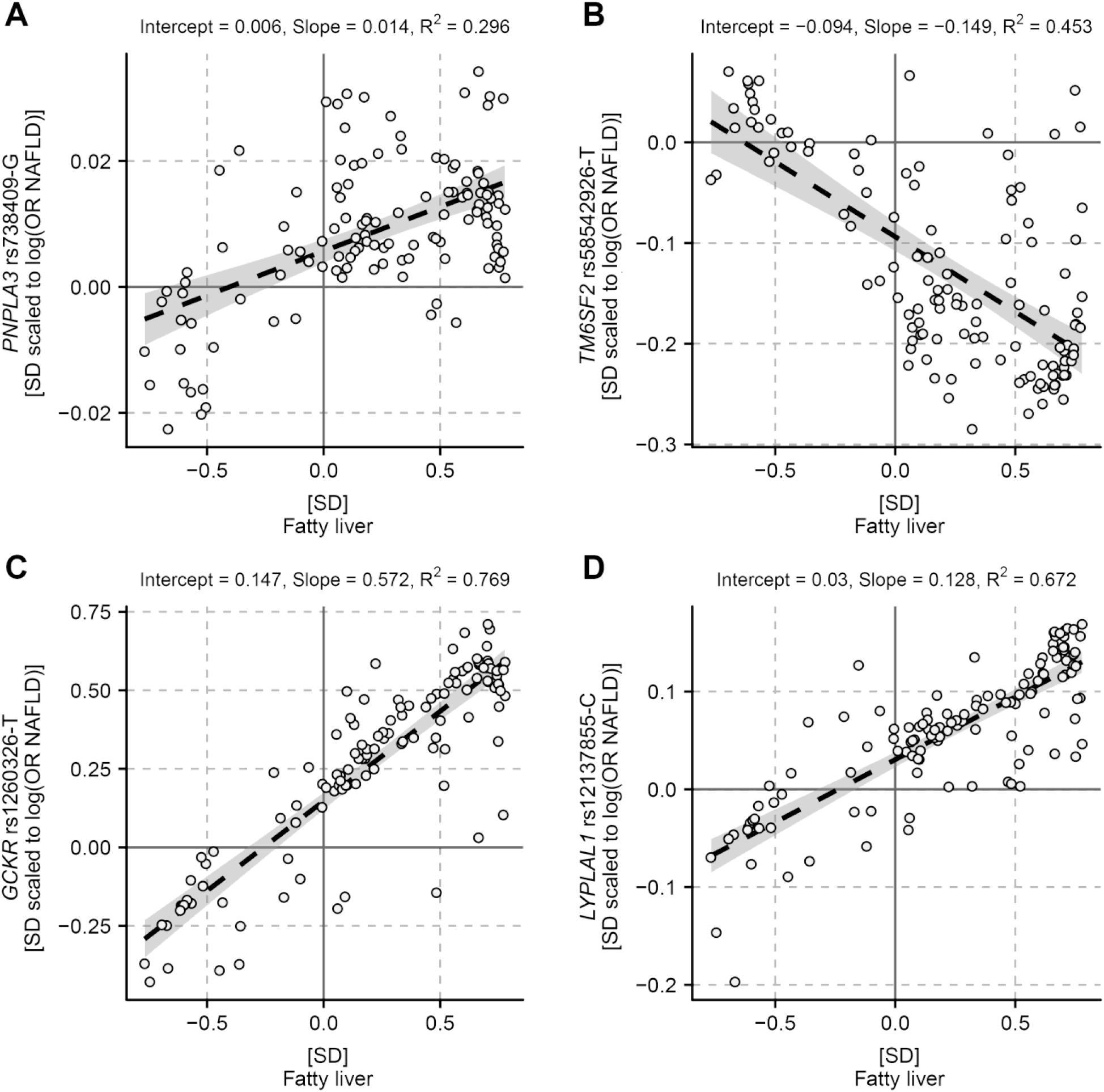
The overall match between the metabolic effects of the NAFLD risk alleles and fatty liver. The dashed line shows the linear fit between metabolic changes associated with fatty liver and *PNPLA3* rs738409-G (A), *TM6SF2* rs58542926-T (B), *GCKR* rs1260326-T (C), and *LYPLAL1* rs12137855-C (D). The grey area indicates the 95% confidence interval for the line. R^2^ is a measure of goodness of fit.

Pairwise comparisons of the risk allele association profiles indicated that the effects of *GCKR* rs1260326-T and *LYPLAL1* rs12137855-C on circulating metabolites were highly similar (R^2^ = 0.71; Supplementary Figure 3A). The overall pattern of metabolic effects of *TM6SF2* rs58542926-T correlated inversely with effects of both *GCKR* rs1260326-T and *LYPLAL1* rs12137855-C (R^2^ = 0.66, Supplementary Figure 3C, and R^2^ = 0.50, Supplementary Figure 3E, respectively). The remaining correlations were weaker (0.19 ≤ R^2^ ≤ 0.24; Supplementary Figure 3).

## Discussion

We assessed fatty liver related metabolic changes in 1,810 young and middle-aged adults from a Finnish population cohort using 123 circulating metabolic measures covering a wide range of metabolic pathways, and further compared the cross-sectional observations with metabolic association profiles of known NAFLD risk alleles obtained from a publicly available metabolomics GWAS including up to 24,925 individuals (21). The studied NAFLD risk alleles resulted in divergent metabolic association profiles. Despite *PNPLA3* rs738409-G being the strongest genetic risk factor for NAFLD (17), it showed a null effect on circulating lipids and metabolites. Association profile of *GCKR* rs1260326-T showed similarities to the cross-sectional fatty liver associations whereas the *TM6SF2* rs58542926-T provided strong statistical evidence to the opposite direction. The present results provide molecular evidence supportive to the recent findings about worsened metabolic features seen in association obesity linked NAFLD but not with “genetic NAFLD”, as defined by NAFLD arising due to risk alleles in *PNPLA3* and *TM6SF2* (19, 27–29).

The differing metabolic effects reflect the biological functions of the risk alleles and suggest that fatty liver can arise from at least three distinct molecular pathways which have divergent consequences on circulating lipids and metabolites. These pathways are summarized in Figure 3. *Pathway I, excessive hepatic glucose levels and amplified lipogenesis: GCKR* rs1260326-T reduces *GCKR* ability to inhibit glucokinase resulting in enhanced hepatic glucose uptake, reduced fatty acid oxidation and increased lipogenesis (30). Hepatic fatty acids can be converted to triglycerides and distributed to downstream pathways to be stored in hepatic lipid droplets or to be secreted in VLDL particles (31), where they can contribute respectively to development of steatosis or to levels of circulating lipids. In agreement with this, *GCKR* rs1260326-T increases risk of fatty liver (12, 14) and raises concentrations of all the apolipoprotein B containing lipoprotein particles and amount of triglycerides within all the examined lipoprotein subclasses (Panel 2 in Figure 1). *GCKR* rs1260326-T associates also with elevated levels of glycolysis related metabolites and circulating fatty acids, as well as increased fatty acid saturation (Panel 2 in Figure 1) compatible with the enhanced glycolytic and lipogenic activities promoted by this variant (30, 32, 33). Further studies are warranted to understand the association between the *GCKR* variant and aberrations in circulating amino acid concentrations. The *LYPLAL1* rs12137855-C variant has a metabolically similar but statistically less robust effect than the *GCKR* rs1260326-T, as seen in highly correlated effect estimates of the two (Supplementary Figure 3A). This provides supportive evidence for LYPLAL1 functioning in hepatic glucose metabolism as proposed in a study by Ahn *et al.* where they showed that *LYPLAL1* inhibition leads to increase in glucose production in human, rat, and mouse hepatocytes (34).

**Figure 3.**
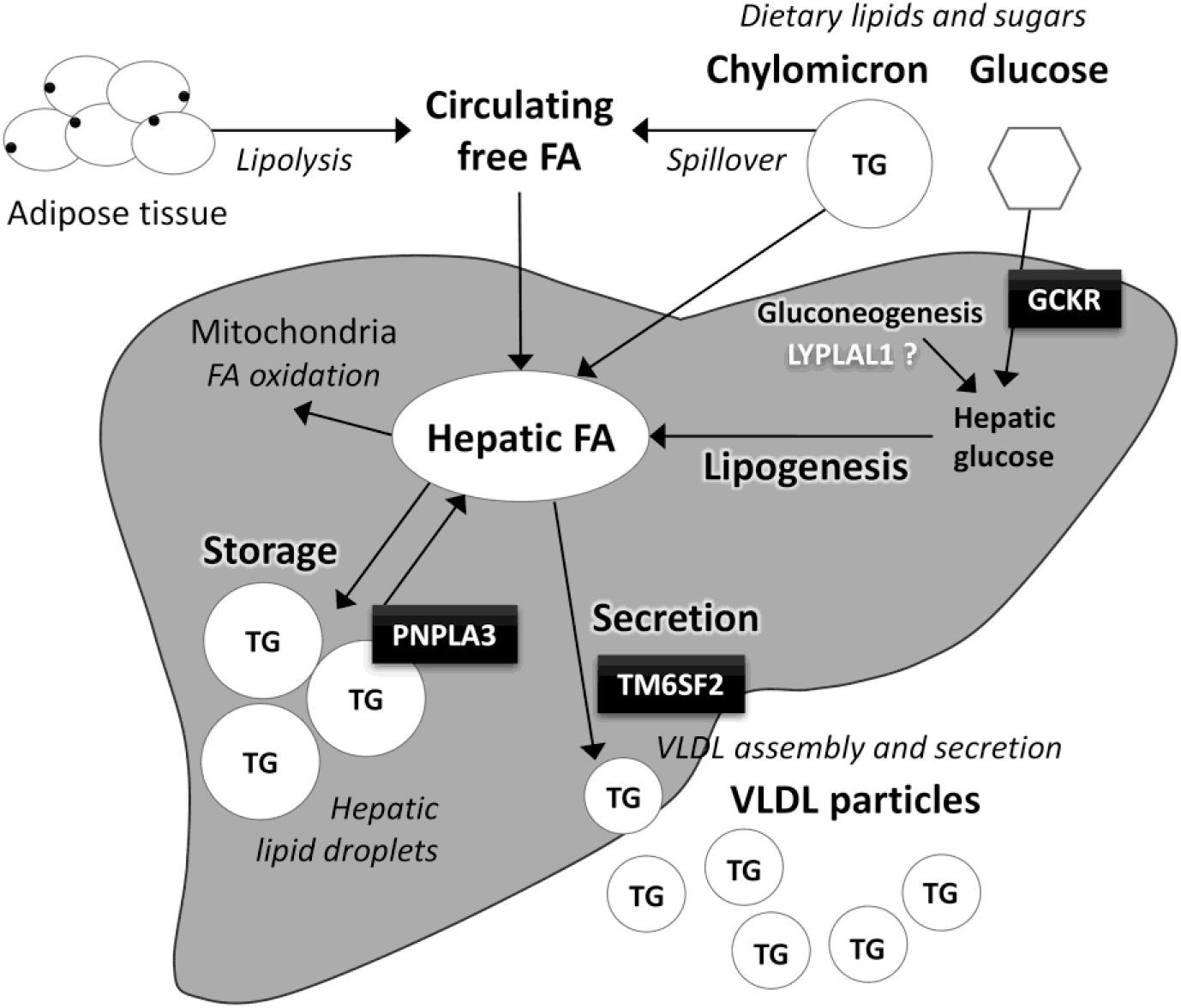
Relation of the studied NAFLD risk alleles to the main pathways in hepatic triglyceride partitioning. Liver converts carbohydrates to lipids in *de novo* lipogenesis. Newly synthesized fatty acids enter to the hepatic fatty acid pool which is also supplied by dietary fats and circulating free fatty acids derived mostly from adipose tissue lipolysis or lipoprotein lipase spillover. Fatty acids can be partitioned to oxidative pathway or esterified to triglycerides that can be stored in hepatic lipid droplets or used for VLDL production to be secreted from the liver. Now, *GCKR* rs1260326-T enhances the lipogenic pathway by providing more substrates for lipogenesis; *LYPLAL1* may be functioning on the same hepatic glucose metabolism and lipogenesis related pathway, as the metabolic effects of *GCKR* rs1260326-T and *LYPLAL1* rs12137855-C are highly similar. Because the hepatic fatty acid pool is located upstream from the storage and secretion pathways, abundance in hepatic fatty acids can contribute to both development of fatty liver and increased production of VLDL. On the contrary, *TM6SF2* rs58542926-T impairs the secretory pathway leading to lipid accumulation into the liver and reduction in levels of circulating lipids and lipoproteins. *PNPLA3* rs738409-G, in turn, enhances triglyceride accumulation to the storage pool by diminishing triglyceride hydrolysis to fatty acids, but does not directly contribute to VLDL secretion, and thus conveys no consequences to circulation.

### Pathway II, reduced VLDL secretion

The TM6SF2 protein contributes to VLDL secretion from the liver (35, 36). In mice, knockdown of *Tm6sf2* causes threefold increase in hepatic triglycerides content while plasma triglycerides and cholesterol are reduced (15). Conversely, overexpression of human *TM6SF2* in mice increases serum lipids while it does not affect liver phenotype (16). The NAFLD risk allele *TM6SF2* rs58542926-T is a loss-of-function variant resulting in a misfolded protein undergoing accelerated degradation (15). Supportive to the previous findings, we observed that this allele associates with reduced concentrations of multiple circulating lipid species while it does not seem to influence on qualitative lipid measures, such as fatty acid saturation (Panel 5 in Figure 1). The inverse associations between *TM6SF2* rs58542926-T and all the circulating VLDL particle subclass concentrations, lipid species within the lipoprotein subclasses, as well as apolipoprotein B concentration suggest that in humans *TM6SF2* rs58542926-T disturbs both lipidation and secretion of VLDL particles. This differs from the observation of a study with *Tm6sf2^-/-^* mice, which provided evidence that mouse Tm6sf2 is required for lipidation of VLDL particles, but lack of it does not influence VLDL secretion (37). The present results are compatible with a study that associated rs58542926 T with a favourable plasma lipid profile (19) and its suggested cardioprotective effect (38).

### Pathway III, impairment of triglyceride mobilisation from hepatic lipid storage

PNPLA3 is localized in the endoplasmic reticulum and lipid droplet membranes in human liver cells (39) and is suggested to be a multifunctional enzyme having both acyltransferase and hydrolase activities on glycerolipids (40–42). NAFLD risk allele rs738409-G enhances triglyceride retention in hepatic lipid droplets by inhibiting their hydrolysis (39, 40, 43) and seemingly has only minimal, if any, contribution to the metabolic traits in circulation (Panel 4 in Figure 1). Our results are in line with other studies showing that *PNPLA3* rs738409-G does not show observable associations with alterations in plasma triglycerides, total cholesterol, HDL-C, LDL-C, nor glucose homeostasis (10, 11), and the present findings extend the same perception to fatty acids, amino acids, and detailed lipoprotein subclass measures. Moreover, our findings are compatible with a mouse model where *Pnpla3*^148M/M^ knock-in mice show no differences in levels of circulating lipids and glucose in comparison to wild-type mice regardless the increase in liver triglycerides (44). These findings advocate that hepatic lipid accumulation can be neutral for circulatory changes and that the strong cross-sectional associations are likely due to lifestyle-related aspects such as dietary factors, excess energy intake or sedentary lifestyle. Dietary lipids and sugars supply the fatty acid pool upstream from the lipid storage and secretion pathways (31), and consequently overnutrition could contribute to both fatty liver development as well as altered metabolic profile.

This study has some limitations. We examine only a limited number of NAFLD risk alleles while there are multiple other genetic pathways contributing to pathogenesis of NAFLD (17, 45). However, the variants studied here are important determinants of liver fat content (13–15, 17, 18) and therefore the current study setting covers some of the fundamental pathways involved in NAFLD pathogenesis. Regarding the cross-sectional associations, ultrasound lacks sensitivity to detect mild steatosis (46) which reduces our power to determine observational associations thus leading to conservative association magnitudes. In addition, ultrasound cannot distinguish simple steatosis from steatohepatitis and therefore we are unable to characterize the liver phenotype in more detail. Alternative techniques for metabolic profiling, such as a more detailed lipidomics platform (47), may reveal other systemic metabolic biomarker changes that could be associated with steatosis induced by the NAFLD risk alleles showing no association to the metabolic traits studied here. However, the metabolic measures captured with the current panel show strong associations with fatty liver, and the studied NAFLD risk alleles show divergent association profiles on the corresponding lipid and metabolite levels informing about the heterogeneous nature of fatty liver.

The present study demonstrates that detailed metabolic profiling can provide extensive information on the biological functions of disease-associated genetic variants, and illustrates how omics data can be utilized in evaluation of molecular mechanisms complex traits, such as NAFLD. The divergence in the direction of the genetic association profiles emphasises that NAFLD is a heterogeneous condition and that its impact on circulating metabolic traits varies depending on the pathogenic mechanism. We highlight the barely discernible metabolic consequence of the strongest genetic determinant of NAFLD, *PNPLA3* rs738409-G, and the cardioprotective metabolic association profile of the *TM6SF2* rs58542926-T. Our findings suggest that hepatic triglyceride accumulation by itself does not necessarily cause metabolic changes increasing the risk of cardiometabolic complications.

Author names in bold designate shared co-first authorship.

## Supporting information

**Supplementary Figure 1. Correspondence of the metabolic effects of *GCKR* rs1260326-T with *GCKR* rs780094-T, and *TM6SF2* rs58542926-T with *NCAN* rs2228603-T.** The dashed line shows the highly matching metabolic effects between the genotypes at the two loci associated with NAFLD risk.

**Supplementary Figure 2. Cross-sectional associations of fatty liver with all the studied 123 metabolic traits, and the corresponding associations of the studied NAFLD risk alleles.** The metabolic traits were adjusted for age, sex, and ten first genetic principal components prior to analysis. Effect estimates of *GCKR* rs1260326-T, *GCKR* rs780094-T, *LYPLAL1* rs12137855-C, *PNPLA3* rs738409-G, *TM6SF2* rs58542926-T, and *NCAN* rs2228603-T on metabolic traits were acquired from a publicly available metabolomics GWAS (17). Biomarker abbreviations as in Supplementary Table 1.

**Supplementary Figure 3. The overall match between the metabolic effects of the NAFLD risk alleles *GCKR* rs1260326-T, *LYPLAL1* rs12137855-C, *PNPLA3* rs738409-G, and *TM6SF2* rs58542926-T.** The dashed line shows the linear fit between the studied risk alleles, and the grey area indicates the 95% confidence interval for the line.

**Supplementary Table 1. NMR based metabolic traits.**

**Supplementary Table 2. Fatty liver associations with circulating metabolic traits.**

**Supplementary Table 3. NAFLD risk allele associations with circulating metabolic traits.**

## References

1. Cohen JC, Horton JD, Hobbs HH. Human fatty liver disease: Old questions and new insights. Science 2011; 332(6037): 1519–1523.

2. Adiels M, Taskinen MR, Packard C, Caslake MJ, Soro-Paavonen A, Westerbacka J, Vehkavaara S, et al. Overproduction of large VLDL particles is driven by increased liver fat content in man. Diabetologia 2006; 49(4): 755–765.

3. Chatrath H, Vuppalanchi R, Chalasani N. Dyslipidemia in patients with nonalcoholic fatty liver disease. Semin Liver Dis 2012; 32(1): 22–29.

4. Kaikkonen JE, Wurtz P, Suomela E, Lehtovirta M, Kangas AJ, Jula A, Mikkila V, et al. Metabolic profiling of fatty liver in young and middle-aged adults: Cross-sectional and prospective analyses of the young finns study. Hepatology 2017; 65(2): 491–500.

5. Mannisto VT, Simonen M, Hyysalo J, Soininen P, Kangas AJ, Kaminska D, Matte AK, et al. Ketone body production is differentially altered in steatosis and non-alcoholic steatohepatitis in obese humans. Liver Int 2015; 35(7): 1853–1861.

6. Mannisto VT, Simonen M, Soininen P, Tiainen M, Kangas AJ, Kaminska D, Venesmaa S, et al. Lipoprotein subclass metabolism in nonalcoholic steatohepatitis. J Lipid Res 2014; 55(12): 2676–2684.

7. Dentin R, Benhamed F, Hainault I, Fauveau V, Foufelle F, Dyck JR, Girard J, et al. Liver-specific inhibition of ChREBP improves hepatic steatosis and insulin resistance in ob/ob mice. Diabetes 2006; 55(8): 2159–2170.

8. Savage DB, Choi CS, Samuel VT, Liu ZX, Zhang D, Wang A, Zhang XM, et al. Reversal of diet-induced hepatic steatosis and hepatic insulin resistance by antisense oligonucleotide inhibitors of acetyl-CoA carboxylases 1 and 2. J Clin Invest 2006; 116(3): 817–824.

9. Seppala-Lindroos A, Vehkavaara S, Hakkinen AM, Goto T, Westerbacka J, Sovijarvi A, Halavaara J, et al. Fat accumulation in the liver is associated with defects in insulin suppression of glucose production and serum free fatty acids independent of obesity in normal men. J Clin Endocrinol Metab 2002; 87(7): 3023–3028.

10. Speliotes EK, Butler JL, Palmer CD, Voight BF, Giant Consortium, MIGen Consortium, Nash CRN, et al. PNPLA3 variants specifically confer increased risk for histologic nonalcoholic fatty liver disease but not metabolic disease. Hepatology 2010; 52(3): 904–912.

11. Romeo S, Kozlitina J, Xing C, Pertsemlidis A, Cox D, Pennacchio LA, Boerwinkle E, et al. Genetic variation in PNPLA3 confers susceptibility to nonalcoholic fatty liver disease. Nat Genet 2008; 40(12): 1461–1465.

12. Speliotes EK, Yerges-Armstrong LM, Wu J, Hernaez R, Kim LJ, Palmer CD, Gudnason V, et al. Genome-wide association analysis identifies variants associated with nonalcoholic fatty liver disease that have distinct effects on metabolic traits. PLoS Genet 2011; 7(3): e1001324.

13. Leon-Mimila P, Vega-Badillo J, Gutierrez-Vidal R, Villamil-Ramirez H, Villareal-Molina T, Larrieta-Carrasco E, Lopez-Contreras BE, et al. A genetic risk score is associated with hepatic triglyceride content and non-alcoholic steatohepatitis in mexicans with morbid obesity. Exp Mol Pathol 2015; 98(2): 178–183.

14. Santoro N, Zhang CK, Zhao H, Pakstis AJ, Kim G, Kursawe R, Dykas DJ, et al. Variant in the glucokinase regulatory protein (GCKR) gene is associated with fatty liver in obese children and adolescents. Hepatology 2012; 55(3): 781–789.

15. Kozlitina J, Smagris E, Stender S, Nordestgaard BG, Zhou HH, Tybjaerg-Hansen A, Vogt TF, et al. Exome-wide association study identifies a TM6SF2 variant that confers susceptibility to nonalcoholic fatty liver disease. Nat Genet 2014; 46(4): 352–356.

16. Holmen OL, Zhang H, Fan Y, Hovelson DH, Schmidt EM, Zhou W, Guo Y, et al. Systematic evaluation of coding variation identifies a candidate causal variant in TM6SF2 influencing total cholesterol and myocardial infarction risk. Nat Genet 2014; 46(4): 345–351.

17. Anstee QM, Day CP. The genetics of NAFLD. Nat Rev Gastroenterol Hepatol 2013; 10(11): 645–655.

18. Sookoian S, Pirola CJ. Meta-analysis of the influence of I148M variant of patatin-like phospholipase domain containing 3 gene (PNPLA3) on the susceptibility and histological severity of nonalcoholic fatty liver disease. Hepatology 2011; 53(6): 1883–1894.

19. Goffredo M, Caprio S, Feldstein AE, D’Adamo E, Shaw MM, Pierpont B, Savoye M, et al. Role of TM6SF2 rs58542926 in the pathogenesis of nonalcoholic pediatric fatty liver disease: A multiethnic study. Hepatology 2016; 63(1): 117–125.

20. Dongiovanni P, Romeo S, Valenti L. Genetic factors in the pathogenesis of nonalcoholic fatty liver and steatohepatitis. Biomed Res Int 2015; 2015: 460190.

21. Kettunen J, Demirkan A, Wurtz P, Draisma HH, Haller T, Rawal R, Vaarhorst A, et al. Genome-wide study for circulating metabolites identifies 62 loci and reveals novel systemic effects of LPA. Nat Commun 2016; 7: 11122.

22. Kettunen J, Tukiainen T, Sarin AP, Ortega-Alonso A, Tikkanen E, Lyytikainen LP, Kangas AJ, et al. Genome-wide association study identifies multiple loci influencing human serum metabolite levels. Nat Genet 2012; 44(3): 269–276.

23. Soininen P, Kangas AJ, Wurtz P, Suna T, Ala-Korpela M. Quantitative serum nuclear magnetic resonance metabolomics in cardiovascular epidemiology and genetics. Circ Cardiovasc Genet 2015; 8(1): 192–206.

24. Würtz P, Kangas AJ, Soininen P, Lawlor DA, Davey Smith G, Ala-Korpela M. Quantitative serum NMR metabolomics in large-scale epidemiology: A primer on -omic technology. Am J Epidemiol 2017; 10.1093/aje/kwx016.

25. Wurtz P, Wang Q, Kangas AJ, Richmond RC, Skarp J, Tiainen M, Tynkkynen T, et al. Metabolic signatures of adiposity in young adults: Mendelian randomization analysis and effects of weight change. PLoS Med 2014; 11(12): e1001765.

26. Wurtz P, Wang Q, Soininen P, Kangas AJ, Fatemifar G, Tynkkynen T, Tiainen M, et al. Metabolomic profiling of statin use and genetic inhibition of HMG-CoA reductase. J Am Coll Cardiol 2016; 67(10): 1200–1210.

27. Lallukka S, Sevastianova K, Perttila J, Hakkarainen A, Orho-Melander M, Lundbom N, Olkkonen VM, et al. Adipose tissue is inflamed in NAFLD due to obesity but not in NAFLD due to genetic variation in PNPLA3. Diabetologia 2013; 56(4): 886–892.

28. Hyysalo J, Gopalacharyulu P, Bian H, Hyotylainen T, Leivonen M, Jaser N, Juuti A, et al. Circulating triacylglycerol signatures in nonalcoholic fatty liver disease associated with the I148M variant in PNPLA3 and with obesity. Diabetes 2014; 63(1): 312–322.

29. Petaja EM, Yki-Jarvinen H. Definitions of normal liver fat and the association of insulin sensitivity with acquired and genetic NAFLD-A systematic review. Int J Mol Sci 2016; 17(5): 10.3390/ijms17050633.

30. Beer NL, Tribble ND, McCulloch LJ, Roos C, Johnson PR, Orho-Melander M, Gloyn AL. The P446L variant in GCKR associated with fasting plasma glucose and triglyceride levels exerts its effect through increased glucokinase activity in liver. Hum Mol Genet 2009; 18(21): 4081–4088.

31. Hodson L, Frayn KN. Hepatic fatty acid partitioning. Curr Opin Lipidol 2011; 22(3): 216–224.

32. Rees MG, Wincovitch S, Schultz J, Waterstradt R, Beer NL, Baltrusch S, Collins FS, et al. Cellular characterisation of the GCKR P446L variant associated with type 2 diabetes risk. Diabetologia 2012; 55(1): 114–122.

33. Santoro N, Caprio S, Pierpont B, Van Name M, Savoye M, Parks EJ. Hepatic de novo lipogenesis in obese youth is modulated by a common variant in the GCKR gene. J Clin Endocrinol Metab 2015; 100(8): 1125.

34. Ahn K, Boehm M, Brown MF, Calloway J, Che Y, Chen J, Fennell KF, et al. Discovery of a selective covalent inhibitor of lysophospholipase-like 1 (LYPLAL1) as a tool to evaluate the role of this serine hydrolase in metabolism. ACS Chem Biol 2016; 11(9): 2529–2540.

35. Mahdessian H, Taxiarchis A, Popov S, Silveira A, Franco-Cereceda A, Hamsten A, Eriksson P, et al. TM6SF2 is a regulator of liver fat metabolism influencing triglyceride secretion and hepatic lipid droplet content. Proc Natl Acad Sci U S A 2014; 111(24): 8913–8918.

36. Sookoian S, Castano GO, Scian R, Mallardi P, Fernandez Gianotti T, Burgueno AL, San Martino J, et al. Genetic variation in transmembrane 6 superfamily member 2 and the risk of nonalcoholic fatty liver disease and histological disease severity. Hepatology 2015; 61(2): 515–525.

37. Smagris E, Gilyard S, BasuRay S, Cohen JC, Hobbs HH. Inactivation of Tm6sf2, a gene defective in fatty liver disease, impairs lipidation but not secretion of very low density lipoproteins. J Biol Chem 2016; 291(20): 10659–10676.

38. Pirola CJ, Sookoian S. The dual and opposite role of the TM6SF2-rs58542926 variant in protecting against cardiovascular disease and conferring risk for nonalcoholic fatty liver: A meta-analysis. Hepatology 2015; 62(6): 1742–1756.

39. He S, McPhaul C, Li JZ, Garuti R, Kinch L, Grishin NV, Cohen JC, et al. A sequence variation (I148M) in PNPLA3 associated with nonalcoholic fatty liver disease disrupts triglyceride hydrolysis. J Biol Chem 2010; 285(9): 6706–6715.

40. Huang Y, Cohen JC, Hobbs HH. Expression and characterization of a PNPLA3 protein isoform (I148M) associated with nonalcoholic fatty liver disease. J Biol Chem 2011; 286(43): 37085–37093.

41. Jenkins CM, Mancuso DJ, Yan W, Sims HF, Gibson B, Gross RW. Identification, cloning, expression, and purification of three novel human calcium-independent phospholipase A2 family members possessing triacylglycerol lipase and acylglycerol transacylase activities. J Biol Chem 2004; 279(47): 48968–48975.

42. Kumari M, Schoiswohl G, Chitraju C, Paar M, Cornaciu I, Rangrez AY, Wongsiriroj N, et al. Adiponutrin functions as a nutritionally regulated lysophosphatidic acid acyltransferase. Cell Metab 2012; 15(5): 691–702.

43. Trepo E, Romeo S, Zucman-Rossi J, Nahon P. PNPLA3 gene in liver diseases. J Hepatol 2016; 765(2): 399–412.

44. Smagris E, BasuRay S, Li J, Huang Y, Lai KM, Gromada J, Cohen JC, et al. Pnpla3I148M knockin mice accumulate PNPLA3 on lipid droplets and develop hepatic steatosis. Hepatology 2015; 61(1): 108–118.

45. Hooper AJ, Adams LA, Burnett JR. Genetic determinants of hepatic steatosis in man. J Lipid Res 2011; 52(4): 593–617.

46. Schwenzer NF, Springer F, Schraml C, Stefan N, Machann J, Schick F. Non-invasive assessment and quantification of liver steatosis by ultrasound, computed tomography and magnetic resonance. J Hepatol 2009; 51(3): 433–445.

47. Kotronen A, Seppanen-Laakso T, Westerbacka J, Kiviluoto T, Arola J, Ruskeepaa AL, Yki-Jarvinen H, et al. Comparison of lipid and fatty acid composition of the liver, subcutaneous and intra-abdominal adipose tissue, and serum. Obesity (Silver Spring) 2010; 18(5): 937–944.

